# Tissue specific function of the *Drosophila* 7SK snRNP in controlling motoneuron synaptic growth

**DOI:** 10.1101/2025.04.29.651232

**Authors:** Giriram Mohana, Nastasja Kreim, Jean-Yves Roignant

**Affiliations:** Institute of Molecular Biology, 55128 Mainz, Germany; Bioinformatic core facility, Institute of Molecular Biology (IMB), 55128 Mainz, Germany; Center for Integrative Genomics, Faculty of Biology and Medicine, University of Lausanne, 10015 Lausanne, Switzerland; Institute of Pharmaceutical and Biomedical Sciences, Johannes Gutenberg-University Mainz, 55128 Mainz, Germany

**Keywords:** 7SK, axons, Alazami, Larp7, P-TEFb

## Abstract

The 7SK snRNP is a ribonucleoprotein complex comprising the non-coding RNA 7SK and the associated proteins MePCE, LARP7, and HEXIM. It regulates transcription in higher eukaryotes by sequestering the positive transcription elongation factor (P-TEFb), preventing premature entry of RNA Polymerase II in elongation. Loss of LARP7 in humans causes the Alazami syndrome, marked by restricted growth, impaired movement, and intellectual disability, though the underlying mechanisms remain unclear. In this study, we show that loss of Larp7 or 7SK RNA in *Drosophila* is viable but impairs locomotion and reduces axonal growth at neuromuscular junctions. Larp7 is enriched in specific motoneurons, where it functions autonomously to promote axogenesis. Reducing P-TEFb abundance partially rescues the locomotion and axonal growth defects, indicating that the 7SK complex mediates this function via transcriptional regulation. Transcriptomic analysis of mutant motoneurons revealed that the 7SK complex primarily regulates long genes with high GC content at their promoters. These findings provide new insights into the tissue-specific roles of the 7SK snRNP in transcription and organismal function.

## Introduction

Gene expression is a tightly regulated process essential for cellular development, homeostasis, and responses to environmental signals. A key regulatory step is promoter-proximal pausing, where RNA Polymerase II (Pol II) begins transcription but pauses shortly after entering elongation. This mechanism fine-tunes gene expression, enabling rapid and coordinated transcriptional activation in response to diverse stimuli (Adelman & Lis, 2012; Chen *et al*, 2018; Core & Adelman, 2019; Noe Gonzalez *et al*, 2021). The release of paused Pol II is tightly controlled by regulatory factors, which includes the 7SK snRNP complex among many others (Akhtar *et al*, 2019; Akhtar *et al*, 2021; Nguyen *et al*, 2001; Yang *et al*, 2001).

The 7SK snRNP complex comprises the non-coding nuclear RNA 7SK and associated proteins, including Methylphosphate Capping Enzyme (MePCE), La-related protein 7 (LARP7), and Hexamethylene bis-acetamide-inducible protein (HEXIM). MePCE stabilizes and caps 7SK RNA (Xue *et al*, 2010), while LARP7 ensures the HEXIM proteins remain bound to 7SK RNA (Markert *et al*, 2008). HEXIM1 and HEXIM2 act as scaffolds (Yik *et al*, 2003), sequestering the positive transcription elongation factor b (P-TEFb), a key regulator of transcription elongation consisting of Cyclin-dependent kinase 9 (Cdk9) and Cyclin T (Marshall & Price, 1995; Peng *et al*, 1998; Peterlin & Price, 2006; Price, 2000; Wada *et al*, 1998). By sequestering P-TEFb, the 7SK complex prevents its interaction with paused Pol II, maintaining the polymerase in a poised state near promoter regions.

Beyond transcriptional regulation, the 7SK complex is implicated in RNA processing, including 3′-end formation (Castelo-Branco *et al*, 2013) and alternative splicing (Barboric *et al*, 2009), underscoring its broader roles in RNA metabolism. It is involved in critical biological processes such as cellular differentiation, development, and stress response (Abuhashem *et al*, 2022; Williams *et al*, 2015; Zeitlinger *et al*, 2007). Dysregulation of the 7SK complex has been linked to various pathological conditions, including cancer (He *et al*, 2008), viral infections (Yik *et al*, 2004), and Alazami syndrome (Alazami *et al*, 2012), a disorder characterized by intellectual disability, impaired growth, and locomotion defects. These connections highlight the importance of unraveling the complex’s tissue-specific roles and regulatory networks to better understand its contributions to health and disease.

Despite progress in elucidating the structure and molecular functions of the 7SK complex, key questions remain unanswered, particularly regarding how its loss leads to specific phenotypes. In this study, we uncovered a tissue-specific function of the 7SK snRNP complex in *Drosophila*. We generated loss-of-function mutants for the 7SK RNA and Larp7, finding that while both mutants are viable, they exhibit impaired locomotion. Larp7 is enriched in a subset of motoneurons and functions autonomously within these cells to support proper locomotion. Disruption of the 7SK complex reduced axonal growth at larval neuromuscular junctions, a defect partially rescued by reducing Cdk9 levels, indicating that this function operates at the transcriptional level. Transcriptomic analysis of mutant motoneurons revealed that the loss of the 7SK complex preferentially affects long genes with high GC content at their promoters and those with cis-regulatory elements associated with strong Pol II pausing.

Together, our findings provide new insights into the tissue-specific functions of the 7SK snRNP complex and its role in regulating transcriptional programs essential for proper neuronal and organismal function.

## Results

### Larp7 is enriched in motoneurons and interneurons

To investigate the in vivo function of Larp7, we first examined its expression pattern during development and adulthood. Using the CRISPR/Cas9 system, we inserted an enhanced Green Fluorescent Protein (eGFP) tag at the 5′ end of the Larp7 open reading frame (ORF). The insertion was validated through qPCR on genomic DNA and western blot analysis using an anti-GFP antibody (Fig. EV1). The resulting recombinant flies exhibited no developmental or behavioral abnormalities compared to wild-type flies, indicating that the eGFP tag did not disrupt Larp7 function. eGFP-Larp7 expression was observed across all developmental stages and tissues analyzed (Figs. 1A and EV1). Regardless of cell type, eGFP-Larp7 was exclusively localized to the nucleus, with pronounced enrichment in the nucleolus. This observation aligns with previous reports of Larp7 localization in other organisms (Slomnicki *et al*, 2016). Interestingly, detailed analysis revealed a subset of midline cells in late-stage embryos displaying stronger eGFP-Larp7 staining (Fig. 1A). To identify these cells, we utilized a series of Gal4 drivers known to mark embryonic midline cells. This analysis showed significant overlap of eGFP-Larp7 expression with cells positive for *Tdc2*, *per*, *MzVUM*, and *engrailed* (Fig. 1B,C). These marked cells include motoneurons, while *engrailed*- positive cells also comprise some interneurons (Fig. 1D).

**Figure 1:**
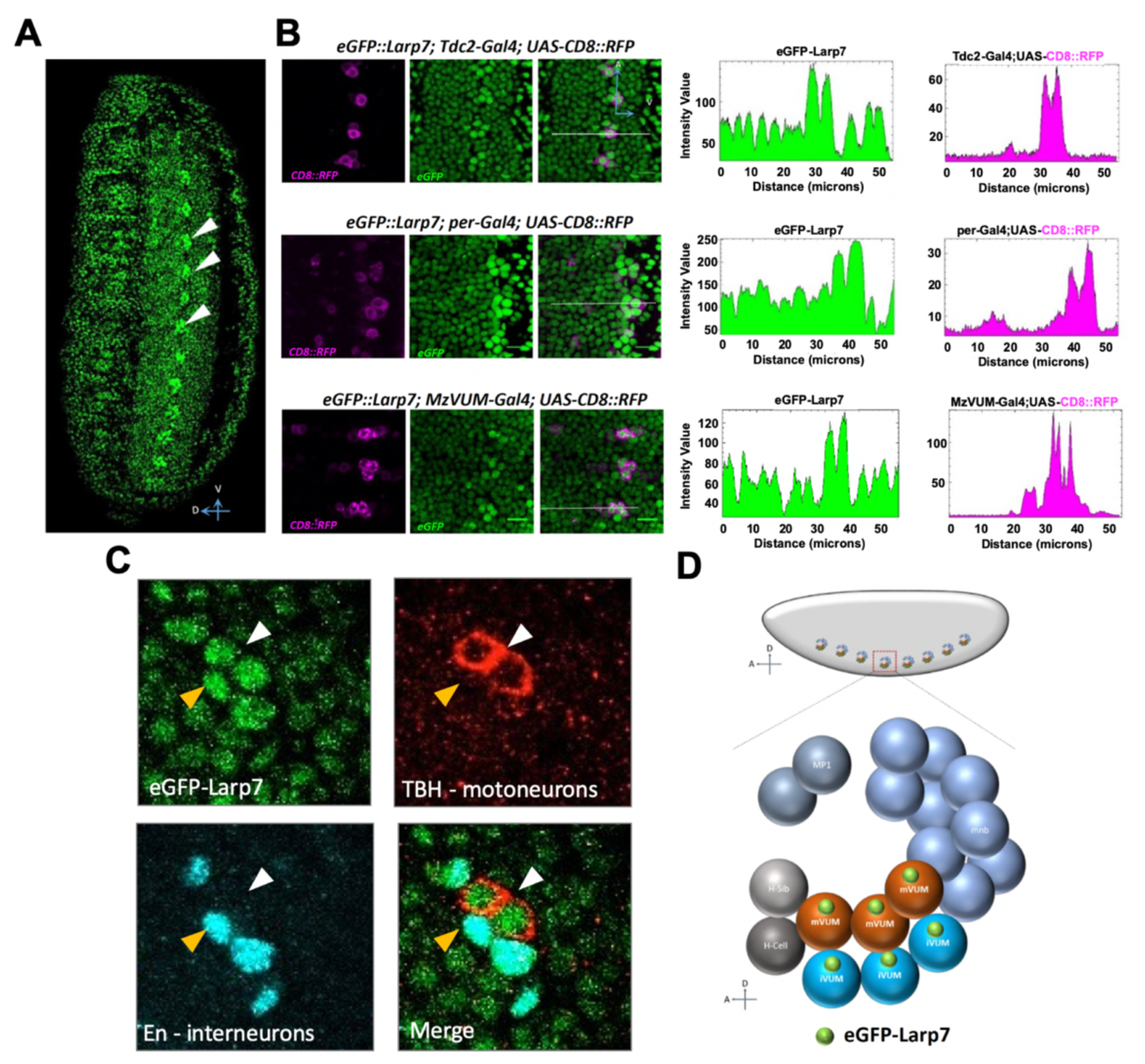
eGFP-Larp7 is enriched at the embryonic midline. **(A)** Representative image of stage 16 embryo. White arrowheads show enriched expression of eGFP-Larp7 in the midline cells. **(B)** Confocal images showing colocalization of eGFP-Larp7 with CD8-RFP driven by indicated Gal4 driver lines. White dotted lines indicate the area analyzed in ImageJ. Graph showing intensity value profile of eGFP-Larp7 and CD8-RFP, Plot-Profile application from ImageJ. **(C)** Confocal images showing colocalization of eGFP-Larp7 (green) with TBH (red) and engrailed (blue). White arrowhead shows enriched expression in motoneurons and yellow arrowhead in interneurons. (**D)** Cartoon depicting the cell types in the midline with eGFP-Larp7 enrichment.

Collectively, these findings demonstrate that Larp7 is ubiquitously expressed but is particularly enriched in a subset of motoneurons and interneurons located at the embryonic midline.

### Larp7 and 7SK RNA are required for locomotion

To further investigate the function of the 7SK complex in *Drosophila*, we generated loss-of-function alleles for the 7SK RNA and *Larp7* using CRISPR/Cas9. For each gene, we created a deletion that removed nearly the entire locus (Fig. EV2A), and these deletions were confirmed via PCR on genomic DNA. As expected, these mutations completely abolished the expression of the respective genes (Fig. EV2B,C). Consistent with previous reports, the loss of Larp7 significantly reduced 7SK RNA stability. In both mutants, the 7SK RNA levels were restored by reintroducing a single copy of either gene (Fig. EV2C). Next, we evaluated the impact of these loss-of-function alleles on *Drosophila* development. Surprisingly, both Larp7 and 7SK RNA mutants were homozygous viable and displayed no developmental delays. However, their lifespans were moderately reduced, with the defect being more pronounced in *Larp7* mutants and double-knockout (DKO) flies (Fig. EV3A). This observation suggests that Larp7 might have additional functions independent of the 7SK snRNP.

Despite the absence of obvious morphological abnormalities, we observed reduced locomotion in single-KO and DKO flies. This locomotion impairment was evident as early as the third larval instar stage (Fig. 2A). Representative traces of peristaltic movements over two minutes are shown in Fig. 2B. Importantly, the locomotion defect in *Larp7* mutants was rescued by introducing an exogenous copy of the Larp7 cDNA, ruling out off-target effects.

**Figure 2:**
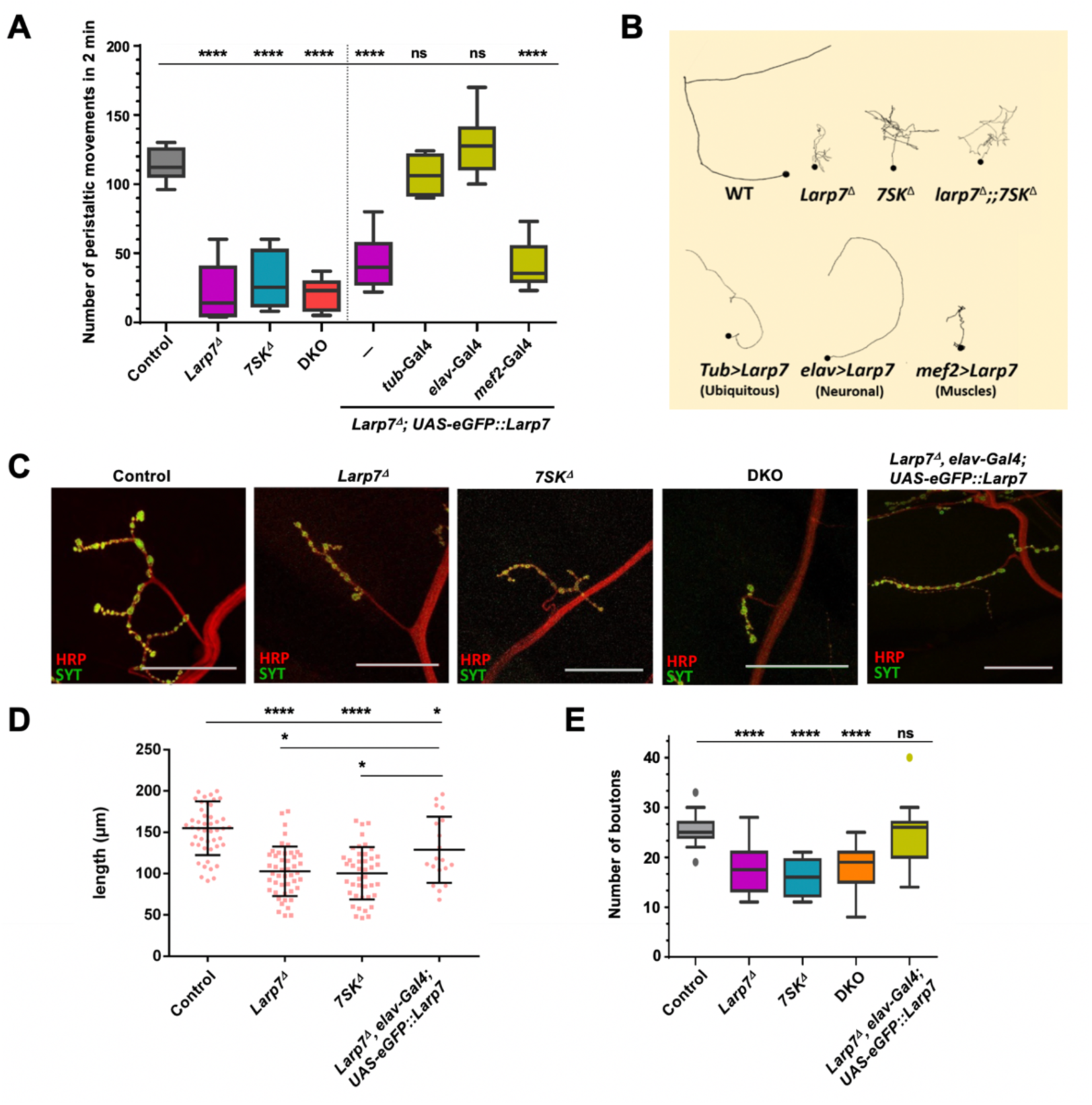
7SK snRNP components are required for larval locomotion and growth of motoneurons. **(A)** Box-plots showing the number of peristaltic movements counted per two minutes for the indicated genotypes. **(B)** Representative tracks of L3 larval movement recorded for 2 minutes. Black dots indicate the position of the larvae at time zero (start point). **(C)** Confocal images of immunostained muscle 4 NMJ for the indicated genotypes. Anti-Synaptotagmin (green) marks the presynaptic terminal of NMJ. Anti-HRP (red) stains the nerve. (scale bar =50uM). **(D)** Quantification of axon growth at the NMJ of muscle 4. **(E)** Quantification of bouton numbers at the NMJ of muscle 4. Control n=19, *Larp7^Δ^* n=20, *7SK^11^* n=16, DKO n=11, *Larp7^Δ^*, elav-Gal4; UAS-eGFP::Larp7 n=11, *tub*-Gal4 n= 10, *elav*-Gal4 n=10, *mef2*-Gal4 n=12. Unpaired t test with Welch’s correction

We hypothesized that the locomotion defects might result from dysfunction in neurons, muscles, or both. To test this, we expressed an exogenous copy of Larp7 in specific tissues of *Larp7* mutants and assessed larval locomotion behavior. Pan-neuronal expression of Larp7 using the *elav-*Gal4 driver fully restored locomotion, whereas expression in muscles using the *mef2-*Gal4 driver had no effect (Fig. 2A,B). These findings highlight the critical role of Larp7 in neuronal tissues for regulating locomotion activity. Similar rescue experiments could not be performed for 7SK RNA mutants because 7SK is expressed via RNA Polymerase III, which is incompatible with the UAS-Gal4 binary expression system.

### Larp7 and *7SK* RNA are required for the growth of motoneurons

The observation that Larp7 controls locomotion through a neuronal function and is enriched in motoneurons led us to investigate the morphology of neuromuscular junctions (NMJs). NMJs serve as the interface between motoneuron axon terminals and muscle fibers, where synaptic activation regulates muscle excitation and, consequently, larval locomotion. NMJs at muscles 4 and 6/7 are frequently studied due to their accessibility and the consistent number of synapses in corresponding body segments among larvae at the same developmental stage. Accordingly, we analyzed the NMJs of muscle 4 in segments A2 and A3 in wandering third-instar (L3) larvae. Notably, we found a significant reduction in axon length in both 7SK and *Larp7* mutants (Fig. 2C,D). Additionally, both mutants exhibited a striking decrease in the number of synaptic boutons (Fig. 2E). The extent of synaptic reduction in DKO larvae was comparable to that observed in each single KO, suggesting that Larp7 and 7SK RNA function together to regulate axonal growth at the NMJ. Importantly, ectopic neuronal expression of Larp7 was sufficient to restore NMJ morphology, underscoring the specificity of the phenotype and the critical role of Larp7 activity in neurons (Fig. 2C-E).

Together, these findings demonstrate that both Larp7 and 7SK RNA are essential for proper axonal growth of motoneurons and the maintenance of NMJ architecture.

### Larp7 acts cell-autonomously in motoneurons

To determine whether Larp7 exerts its function in a cell-autonomous manner, we ectopically expressed Larp7 in various neuronal populations within the *Larp7* mutant background and examined its effect on larval locomotion. Expression of Larp7 in neuroblasts using the *Insc-*Gal4 driver did not rescue locomotion, suggesting that Larp7 is dispensable in this cell type (Fig. 3A). Similarly, expressing Larp7 in glial cells with the *repo-*Gal4 driver produced larvae with locomotion defects comparable to the *Larp7* mutant alone. However, several motoneuron-specific Gal4 drivers partially or fully rescued the locomotion phenotype, indicating that Larp7 likely functions cell-autonomously in motoneurons.

**Figure 3:**
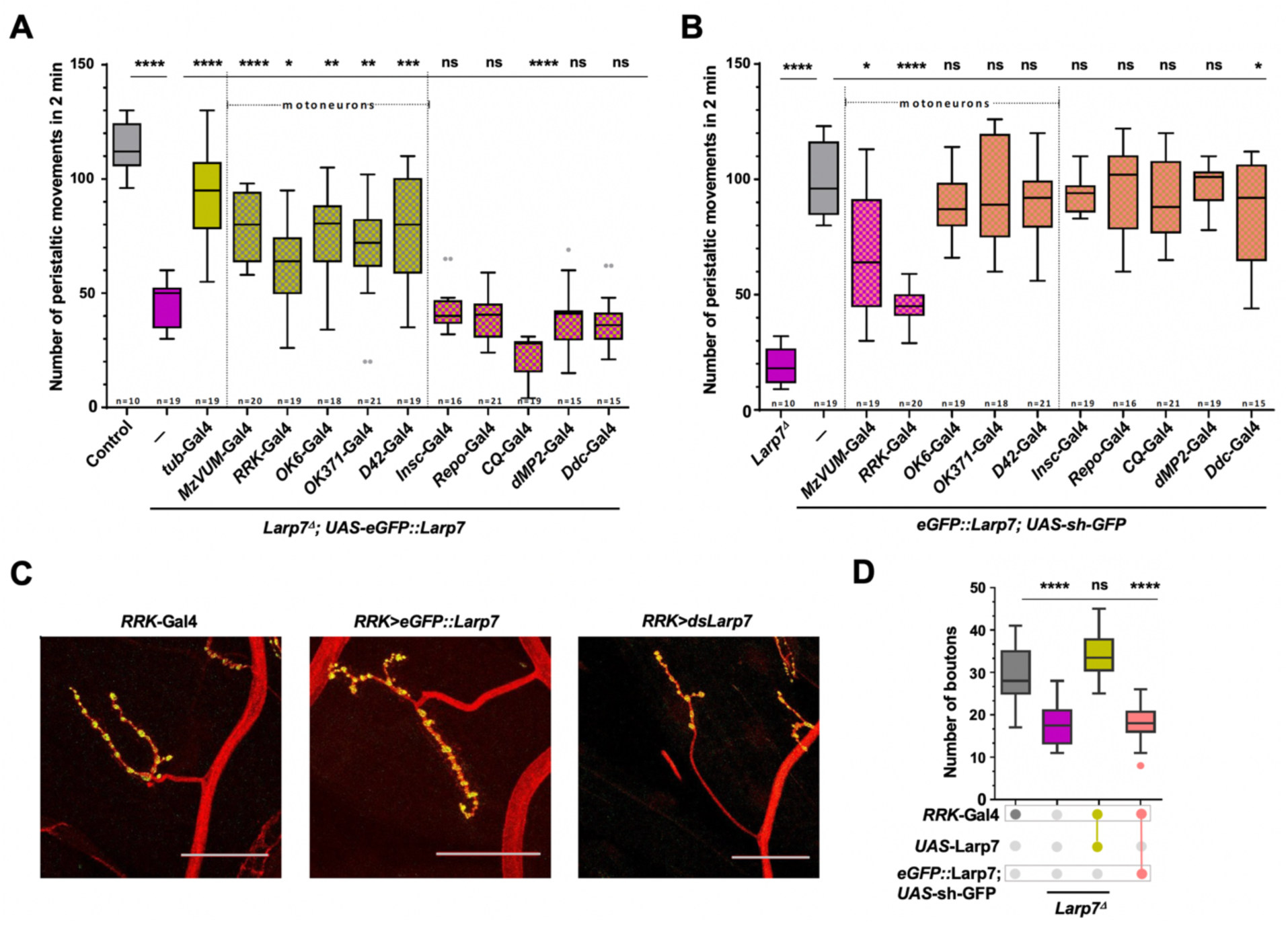
Motoneurons require 7SK snRNP components for locomotion and axonal growth. **(A)** Quantification of the crawling assay upon overexpression of Larp7 using indicated Gal4 drivers. **(B)** Quantification of the crawling assay upon Larp7 knock down using indicated drivers. **(C)** Representative images of NMJ at muscle 4 of the indicated genotypes. Scale bars = 50uM. **(D)** Quantification of bouton numbers in the muscle 4 NMJ. *RRK*-Gal4 n=16, *Larp7^Δ^* n=20, *RRK*-Gal4, UAS-Larp7 n=20, *RRK*-Gal4: eGFP::Larp7; UAS-sh-GFP n=24. Unpaired t test with Welch’s correction.

To confirm this, we specifically depleted Larp7 in different neuronal cell types (Fig. 3B) using the in vivo GFP interference (iGFPi) method as described in (Neumuller *et al*, 2012; Pastor-Pareja & Xu, 2011). iGFPi employs double-stranded RNA (dsRNA) targeting GFP to knock down (KD) GFP-tagged fusion proteins. In this study, we used iGFPi to deplete eGFP-Larp7. A GFP dsRNA line that achieved the most efficient KD (Fig. EV4) was crossed with flies carrying the same Gal4 drivers used in the overexpression experiments. Consistent with the rescue data, depletion of Larp7 in motoneurons caused mild to severe locomotion defects in third-instar larvae, while depletion in other neuronal cell types had no effect (Fig. 3B). Notably, depletion with the motoneuron-specific driver *RRK-*Gal4—which targets the aCC and RP2 motoneurons—produced the strongest defects. Among these, RP2 motoneurons are responsible for innervating muscle 4 in each hemisegment. Other motoneuron drivers, such as *OK6-*Gal4 and *OK371-* Gal4, are expressed in all motoneurons, but their impact was less pronounced than that of *RRK-*Gal4. We attribute this variability to differences in the expression strength of individual Gal4 drivers, which may affect the KD efficiency.

Next, we investigated whether the growth of NMJs at muscle 4 was also affected in a cell-autonomous manner. Consistent with the locomotion results, expressing Larp7 using the *RRK-*Gal4 driver rescued the NMJ growth defects, while depletion of Larp7 with the same driver induced growth impairments (Fig. 3C,D).

Together, these findings demonstrate that Larp7 functions cell-autonomously in motoneurons to regulate their growth and ensure proper larval locomotion.

### The locomotion and axonal growth defects result from P-TEFb hyperactivation

The best-characterized function of the 7SK snRNP is the regulation of transcriptional pausing through sequestration of P-TEFb. Based on this, we hypothesized that the locomotion phenotype and axonal growth defects observed in the 7SK mutants might result from transcriptional dysregulation due to hyperactivation of P-TEFb. If this were the case, reducing P-TEFb levels should ameliorate the defects associated with the loss of 7SK snRNP function. To test this hypothesis, we knocked down Cdk9 in the *Larp7* mutant background using *elav-*Gal4 and *RRK-*Gal4. Strikingly, we found that Cdk9 KD in all neurons, as well as in motoneurons, partially rescued the crawling defects in *Larp7* mutants (Fig. 4A). Even more remarkably, we observed complete rescue of the number of synaptic boutons and axonal length at the NMJ of muscle 4 in these larvae (Fig. 4B,C). Similar results were obtained in the double KD of Larp7 and Cdk9 (Fig. 4A-C). Importantly, Cdk9 KD alone in wild-type neurons is embryonic lethal, underscoring the tightly regulated nature of Cdk9 levels in neurons and the essential role of Larp7 and the 7SK snRNP in this regulation.

**Figure 4:**
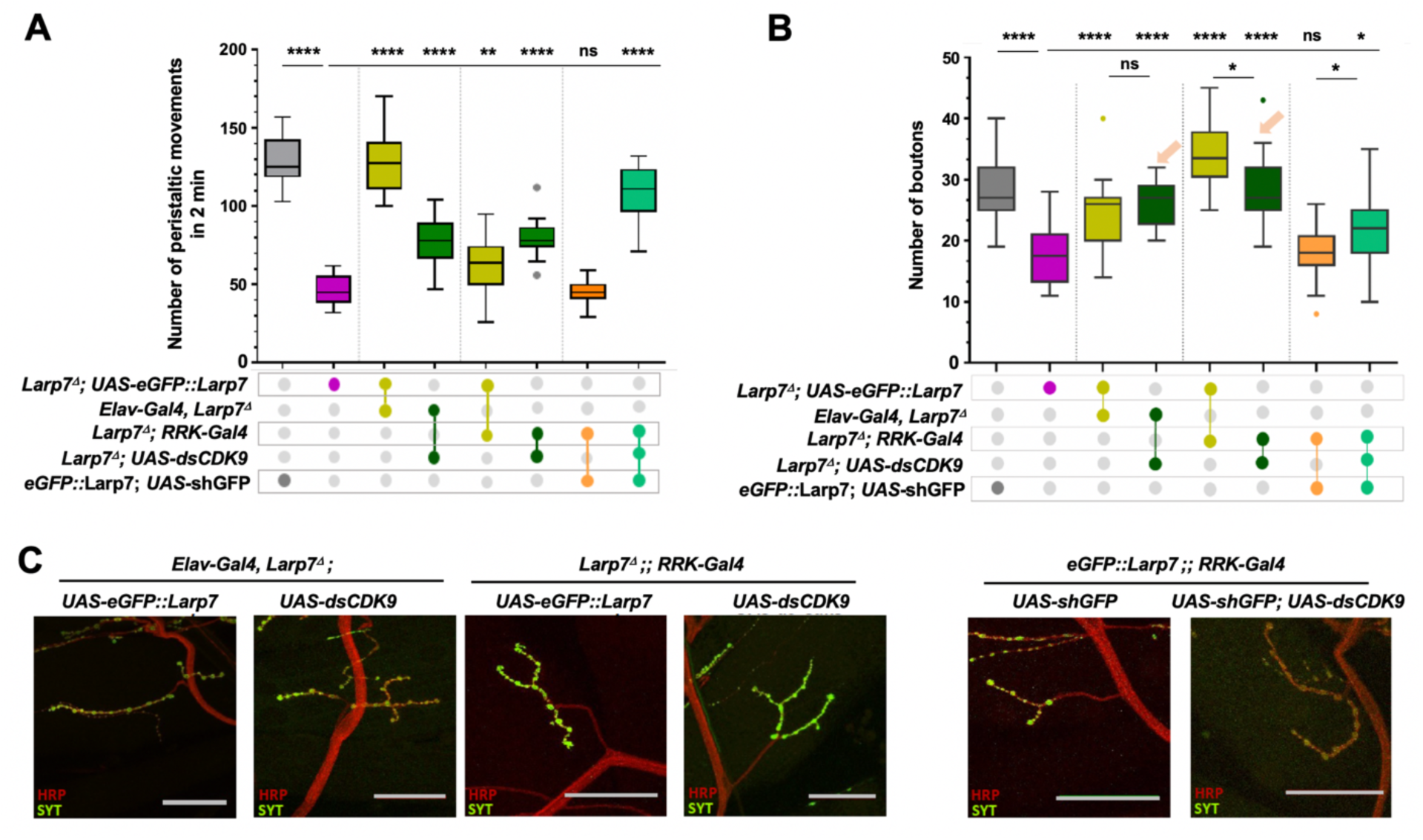
Reducing Cdk9 levels rescues Larp7 depletion defects. **(A)** Crawling assay showing partial rescue of *Larp7* mutant defect upon Cdk9 KD in all neurons (*elav*- Gal4) or in motoneurons (*RRK*-Gal4). N comprises between 10 and 20. Unpaired t test with Welch’s correction. **(B)** Quantification of bouton numbers at the NMJ of muscle 4. N comprises between 15 and 23. Unpaired t test with Welch’s correction. **(C)** Representative confocal images of muscle 4 NMJ of the indicated genotypes.

Collectively, our findings strongly suggest that the 7SK snRNP controls locomotion and axonal growth at the NMJ through the regulation of P-TEFb.

### Transcriptional regulation by the 7SK RNP complex in motoneurons

Since our results suggest transcriptional regulation in motoneurons by components of the 7SK snRNP complex, we next investigated the transcriptome of aCC and RP2 cells, the motoneurons most sensitive to the loss of the 7SK snRNP. Motoneurons from control, *Larp7* KO, and 7SK KO embryos were isolated by fluorescence-activated cell sorting (FACS) at the stage of differentiation (stages 15-16). The identity of the isolated cells was validated by RT-qPCR (Fig. EV5A,B), and the RNA extracts were processed for high-throughput sequencing. Overall, *Larp7* KO RRK-positive cells exhibited more differentially regulated genes (2,998) compared to 7SK KO-positive cells (1,012) (Fig. 5A,B). Among them, 106 up-regulated and 294 down-regulated genes were common to both mutants. We focused on these shared genes for further analysis, as they likely reflect regulation by the entire 7SK snRNP complex. Gene ontology analysis revealed that up-regulated genes are highly enriched for developmental processes (Fig. 5C), while down-regulated genes are associated with chromosome organization, the cell cycle, and metabolic processes (Fig. 5D). These findings suggest that both genes regulate a common set of genes but also have some unique targets—especially Larp7—indicating again an independent function for this protein.

**Figure 5:**
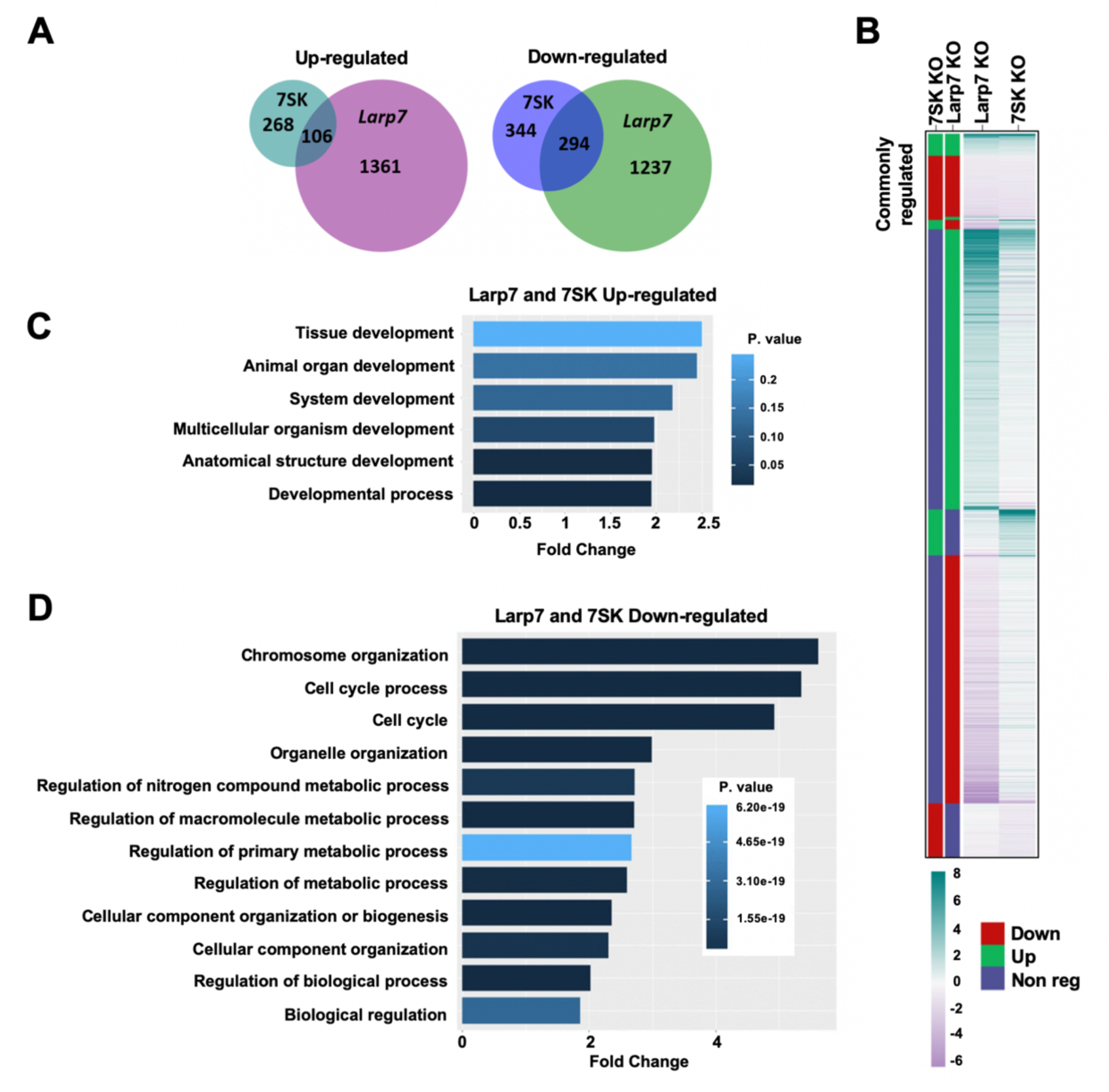
Transcriptomic analysis of 7SK snRNP mutants. **(A)** Euler’s diagram showing the overlap of differentially regulated genes between *7SK* and *Larp7* mutants. **(B)** Heat map of all differentially transcripts in *Larp7* and *7SK* mutant motoneurons. Genes commonly regulated in both mutants are clustered on the top. Diff.reg – indicated the differentially regulated gene with up regulation in green, down regulation in red, and non-regulated in blue. **(C, D)** Gene ontology of commonly up-regulated (C) and down-regulated (D) genes in *Larp7* and *7SK* mutant motoneurons.

Previous studies have shown that the rate of RNA pol II elongation can impact alternative splicing (de la Mata et al., 2003; Khodor et al., 2011; Moehle et al., 2014). Given that the 7SK snRNP complex regulates the recruitment of P-TEFb, we hypothesized that its loss might affect splicing. However, our analysis of splicing events revealed few changes (data not shown), suggesting that the release of pausing or the increase in transcription elongation resulting from 7SK depletion in motoneurons is not sufficient to massively alter alternative splicing.

### The 7SK RNP complex restricts the expression of long genes that are strongly paused

To identify the defining features of snRNP targets, we examined specific gene characteristics among the commonly regulated genes in 7SK and *Larp7* mutants. Several parameters were analyzed, including gene length, GC-rich content, and the presence of specific motifs. The most significant change observed in gene size was the increased total length of up-regulated genes compared to non-regulated and down-regulated genes (Fig. 6A). A similar pattern was found in up-regulated genes in 7SK KO motoneurons, but not in *Larp7* KO. Consistently, the maximum intron size was also significantly larger in the commonly up- regulated genes and in 7SK KO up-regulated genes (Fig. 6B). In contrast, no significant changes were observed in exon length (Fig. 6C).

**Figure 6:**
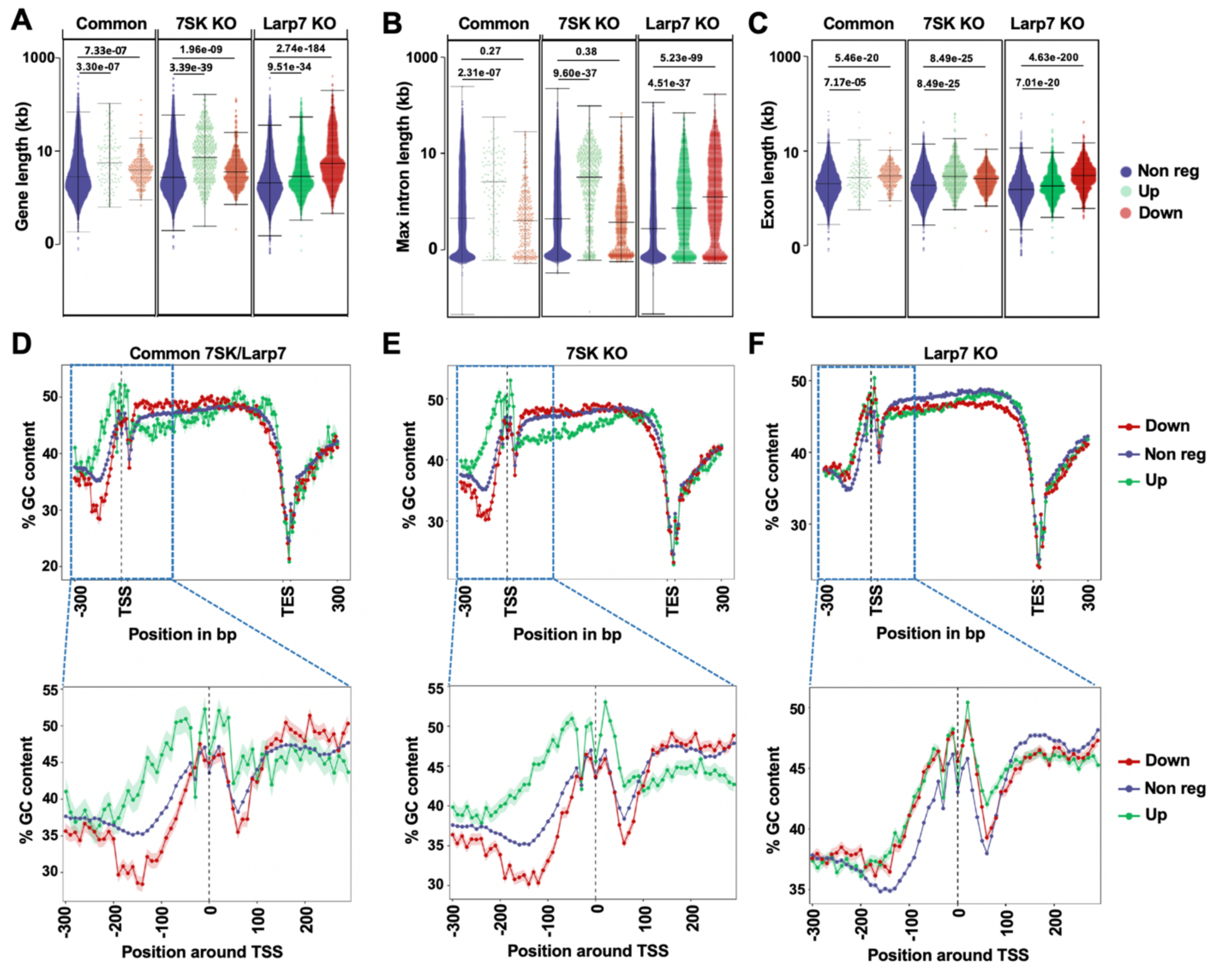
Gene features of 7SK snRNP regulated genes. **(A)** Gene length of differentially regulated genes in the indicated conditions. Up-regulated genes are shown in green, down-regulated genes in red and non-regulated genes in blue. **(B)** Maximum intron length. **(C)** Exon length of aforementioned conditions. **(D-F)** Top: Metagene GC profile of differentially regulated genes in commonly mis-regulated (D), *7SK* (E) and *Larp7* mutants (F). 300bp upstream of transcription start site (TSS) and 300bp downstream of transcription end site (TES) was included in the analysis. Bottom: enlarged region of metagene profiles, 300bp around TSS. Up-regulated genes are indicated in green, down regulated in red and non-regulated in blue.

We next analyzed the GC content of these genes. Notably, we found that commonly up-regulated genes had higher GC content at their promoters, whereas down-regulated genes had lower GC content (Fig. 6D). A similar trend was observed in 7SK-regulated genes, while Larp7-regulated genes did not exhibit this pattern (Fig. 6E,F). We also assessed the GC content within the gene body. Compared to non-regulated genes, both commonly and 7SK up-regulated genes had higher GC content up to 100 bp downstream of the transcription start site (TSS), after which this content substantially decreased within the gene body (Fig. 6D-F). Altogether, this analysis suggests that genes affected by the loss of 7SK snRNP components share distinct features: long genes with high GC content at their promoters are more likely to be up- regulated, while genes with lower GC content at their promoters tend to be down-regulated. Given that RNA polymerase II (Pol II) pausing is associated with GC-rich promoters, these results imply that the 7SK snRNP preferentially restricts the expression of genes that are highly paused.

To validate these findings, we next examined whether the genes up-regulated in the absence of 7SK snRNP function contain characteristic motifs associated with Pol II pausing (Hendrix et al., 2008). For this analysis, we focused on the region spanning +300 bp to -300 bp around the TSS (Fig. 7). Indeed, we found that the promoters of these genes were enriched for motifs such as GAGA, Initiator, DPE, and pause button (Fig. 7). We also observed an enrichment of the TATA motif in differentially regulated genes, although the distribution of this motif differed between up-regulated and down-regulated genes (Fig. 7). The only additional feature identified in down-regulated genes was the enrichment of the DRE motif (Fig. 7).

**Figure 7:**
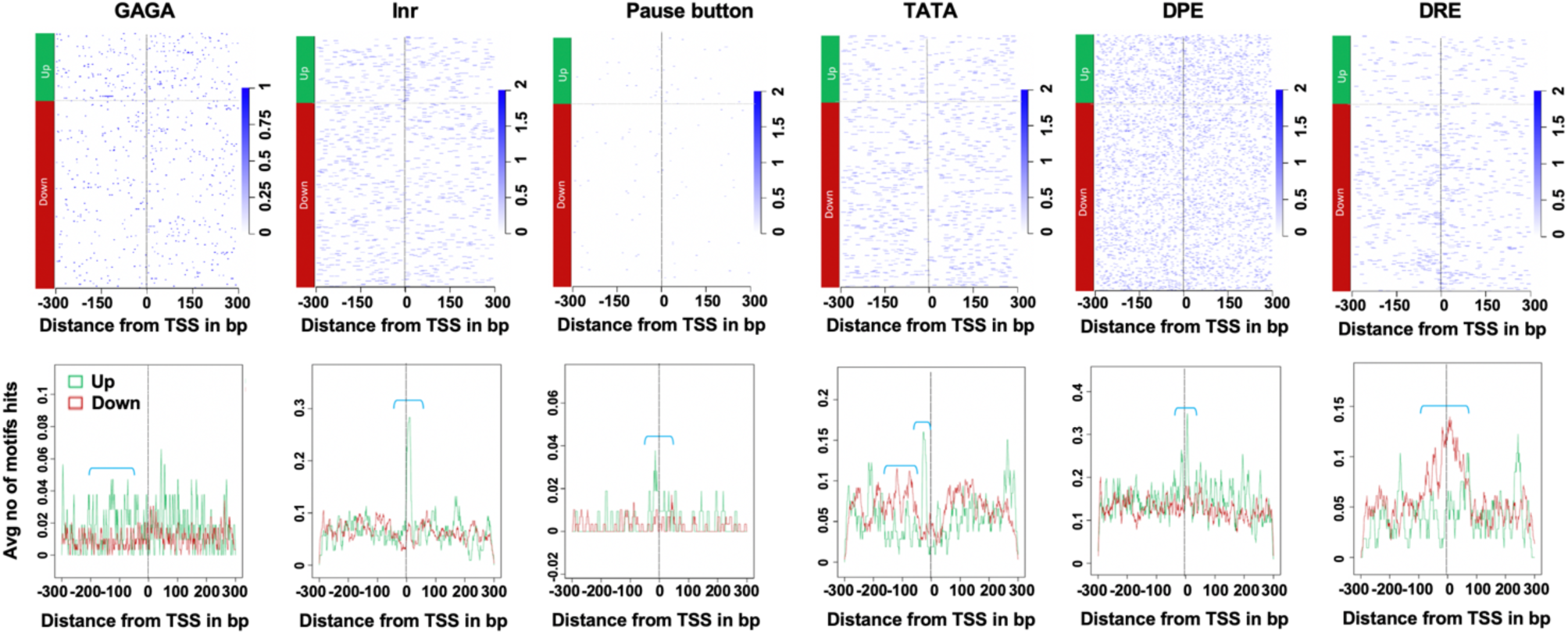
Motif analysis of differentially regulated genes in 7SK snRNP mutant motoneurons. Enrichment of indicated motifs in the commonly regulated genes. Up-regulated genes are shown in green, down-regulated in red and non-regulated in blue. 300bp around TSS was used for the motif enrichment analysis. Downward brackets (turquoise) highlight the enriched region for indicated motifs.

Collectively, these results indicate that genes differentially regulated in 7SK snRNP mutant motoneurons exhibit specific sequence signatures in both their promoters and gene bodies, which likely direct their regulation by the 7SK complex. Notably, genes that show increased expression in the absence of the complex appear to be those that are strongly paused.

### Larp7 function is evolutionarily conserved

Loss of LARP7 in humans leads to Alazami syndrome (AS), a condition characterized by multiple developmental defects. A key feature in all reported AS patients is a significant delay in motor development, suggesting that the 7SK snRNP complex plays an evolutionarily conserved role in the axonal growth of motor neurons. While Drosophila 7SK RNA is approximately 100 nucleotides longer than its vertebrate counterpart and shares less than 50% sequence similarity, the proteins associated with the 7SK RNA, including LARP7, are highly conserved across species. To test whether vertebrate LARP7 is functional in Drosophila, we cloned the human (hLARP7) and Xenopus (xLARP7) LARP7 orthologs, each tagged with GFP. These fusion proteins were expressed in Drosophila S2R+ cells that had been treated with dsRNA targeting the 5’ UTR of Drosophila *Larp7*. As readout, we measured the levels of 7SK RNA by RT-qPCR. Remarkably, we found that both human and Xenopus LARP7 rescued Drosophila 7SK RNA levels to the same extent as Drosophila Larp7 (Fig. 8A). This finding suggests that the molecular function of Larp7 is conserved across evolution.

**Figure 8:**
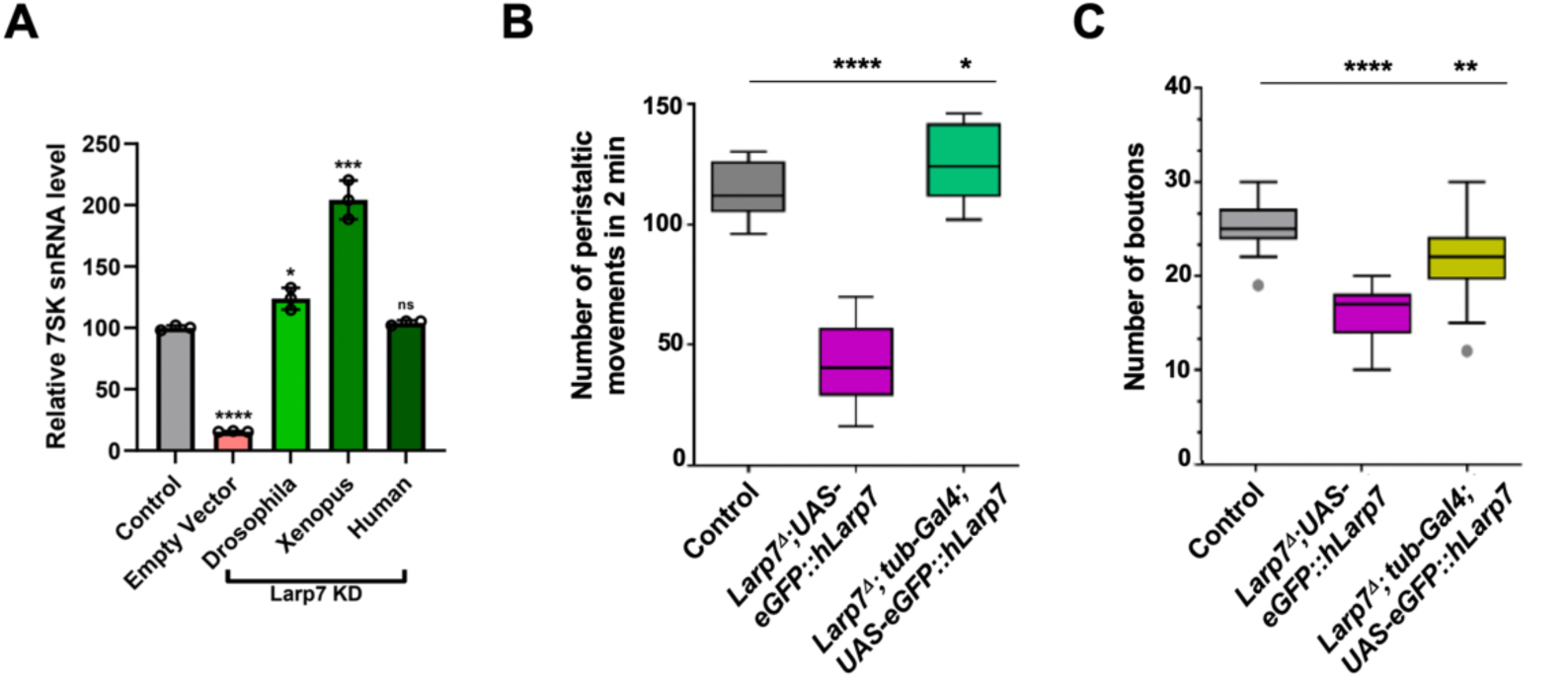
Human LARP7 can rescue *Drosophila Larp7* mutant defects. **(A)** RT-qPCR monitoring the level of *7SK* RNA in *Drosophila Larp7* KD S2R+ cells. cDNA of *dLarp7*, *hLarp7* and *xLarp7* were expressed ectopically with the actin promoter. **(B)** Crawling assay showing the rescue of *Larp7* mutant locomotion defect by hLarp7. N comprises between 10 and 20. Unpaired t test with Welch’s correction. **(C)** Bouton numbers in muscle 4 NMJ of indicated genotypes. Human Larp7 expression in *Drosophila* Larp7 KO larvae rescues the bouton numbers. N comprises between 15 and 25. Unpaired t test with Welch’s correction.

To further investigate functional conservation in vivo, we generated transgenic flies expressing human LARP7 under the control of the UAS promoter. Strikingly, expression of hLARP7 in all tissues using the *tubulin*-Gal4 driver completely rescued the locomotion and axonal growth defects in *Larp7* mutant L3 larvae (Fig. 8B,C). These results demonstrate that Larp7’s function within the 7SK complex is evolutionarily conserved from insects to vertebrates.

## Discussion

In this study, we identified a specific role of the 7SK complex in regulating axonal growth at larval neuromuscular junctions. We discovered that one component of the complex, Larp7, is enriched in a subset of motoneurons and interneurons, where it acts autonomously to promote axonal growth. Loss of 7SK complex function is partially rescued by reducing Cdk9 levels, suggesting that its role in locomotion is mediated through transcriptional activity and release of RNA Pol II pausing. These findings reveal that, despite a predicted general function, the 7SK complex predominantly affects a subset of neuronal cells in vivo. This aligns with observations that patients with mutations in the *LARP7* gene can survive to adulthood but exhibit specific clinical features, including intellectual disability and mobility deficiencies.

The observation that Larp7 is enriched in a subset of motoneurons and interneurons was not anticipated, as well as its specific importance in these cells in vivo. Nevertheless, previous work in mouse cells already hinted for an important role in neurodevelopment. Mouse 7SK RNA level was shown to be enriched in the mouse brain as compared to other tissues, and its knock down impairs neuronal differentiation of embryonic stem cells (Bazi *et al*, 2018). Another study demonstrated a cytoplasmic role for 7SK snRNA in axonal localization of a subset of transcripts (Briese & Sendtner, 2021). Here we show that the role in neurodevelopment is not only conserved in Drosophila but also is largely restricted to this function.

Furthermore, we demonstrate that it worked through the canonical function in transcriptional regulation. Whether a cytoplasmic function also operates in Drosophila is currently unknown. The reason why the 7SK snRNP function is more important in motoneurons is unclear. Increased fine tuning of gene expression in these specialized cells might be more critical. Integration of multiple signals might coordinate nuclear and cytoplasmic functions of the complex, enabling proper neuronal differentiation at the right time.

Promoter proximal pausing is a widespread phenomenon which has been identified as an important regulatory checkpoint that allows the release of Pol II into productive elongation to be tightly controlled (Core & Adelman, 2019; Gaertner & Zeitlinger, 2014). Pol II pausing is prevalent among developmental genes during *Drosophila* embryogenesis (Zeitlinger *et al*., 2007) and allows cells in a tissue to synchronously activate gene expression (Lagha *et al*, 2013). Pausing is influenced through a number of factors including DNA/RNA sequences, nucleosomes, as well as DSIF and NELF, which are phosphorylated by P-TEFb for the release of RNA Pol II into elongation. Consistent with the fact that the 7SK complex controls locomotion through the control of Pol II pausing we have found that upregulated genes in the 7SK loss of function are enriched for developmental processes and contains motifs associated with strong Pol II pausing such as GAGA, Inr and pause button. These motifs serve to recruit transcription factors and chromatin remodeling machineries, opening the chromatin within the promoter regions, which in turn favors stable Pol II binding at promoters along with transcription initiation factors. This stable association has been associated with stronger pausing. Interestingly these genes also tend to be longer and enriched in G/C content at their promoter regions. High G/C content favors the formation of RNA/DNA hybrids, which present an obstacle for Pol II progression. This can lead to Pol II backtracking, which also contributes to longer pausing (Nechaev *et al*, 2010; Sheridan *et al*, 2019). Therefore, our work suggests that the absence of 7SK snRNP specifically stimulates the transcription of a subset of target genes that are highly paused. An alternative scenario is that the 7SK snRNP regulates transcription of all genes but a global feedback stabilization of mRNA may mask the transcriptional changes on less paused genes, as has been suggested earlier (Haimovich *et al*, 2013; Sun *et al*, 2012; Timmers & Tora, 2018). Distinguishing between these possibilities will require the examination of nascent RNA levels by techniques such as pro-seq, which remain challenging in vivo.

How does P-TEFb recognize its targets genes? A number of transcription factors, including c-Myc, P53, the human immunodeficiency virus transactivator Tat or the bromodomain protein 4 (Brd4) were shown to directly bind and target P-TEFb to paused RNA Pol II (AJ *et al*, 2016; Zhou *et al*, 2012). The super elongation complex (SEC) composed of ELL, AFF and ENL/AF9 proteins has also been proposed to deliver and tether P-TEFb to promoters through association with mediator and integrator complexes (Gardini *et al*, 2014; Lu *et al*, 2016; Takahashi *et al*, 2011). In addition, recent data suggest that delivery of P-TEFb to their target genes can be achieved directly by the 7SK complex itself. Indeed fractions of 7SK snRNA and Larp7 were shown to be associated with chromatin, raising the possibility that the 7SK complex may directly interact with P-TEFb dependent genes (D’Orso & Frankel, 2010; Flynn *et al*, 2016; McNamara *et al*, 2013; McNamara *et al*, 2016). Dissociation of chromatin bound 7SK complex in response to specific cellular stimuli was proposed to induce rapid transcriptional activation of primary response gene. This could be achieved through the activity of RNA binding proteins (RBPs) including the splicing factor SRSF2 bound to nascent transcripts, RBM7 as well as the RNA helicase DDX21 (Bugai *et al*, 2019; Calo *et al*, 2015; Ji *et al*, 2013). How does P-TEFb distribute among transcription factors, the SEC, RBPs and how much prior targeting of the 7SK complex to promoters is required for pause release are major questions in the field. Our data show that despite the lack of the 7SK snRNP, free P-TEFb can still stimulate the transcription of genes containing features of pausing implying that there is no prerequisite of targeting the 7SK snRNP to promoters for pause release. This is consistent with previous work in human cells and mouse embryonic stem cells showing that the depletion of 7SK snRNP lead to upregulation of a subset of P-TEFb dependent genes, arguing against a strict requirement for prior anchoring of the 7SK snRNP to chromatin (Castelo-Branco *et al*., 2013; Egloff *et al*, 2017). One possibility is that prior anchoring of the 7SK snRNP might instead be a mean to regulate the dynamic of transcriptional activation of P-TEFb dependent genes rather than their selection. Further studies should address this possibility.

Recent work in Drosophila investigating the function of the Hexim homolog Bin3, demonstrated its involvement in fecundity as well as in fly climbing ability (Palumbo *et al*, 2024). However the mechanism underlying these defects has remained unclear. Based on our data we proposed that Bin3 controls locomotion likely by promoting axonal growth through the sequestration of the P-TEFb complex.

Interestingly *Bin3* loss of function flies also harbor a held-out wing phenotype which is not observed in the *Larp7* and *7SK* mutants, indicating that wing posture is controlled by a different mechanism. This is consistent with the finding that this function depend on a motif in Bin3 that is not required for 7SK binding (Palumbo *et al*., 2024). We note that the molecular consequences of depleting Larp7 is more severe than in the 7SK snRNA KO indicating that Larp7 likely harbors a 7SK snRNP-independent function. Interestingly Larp7 was recently shown to promote ribose methylation of the U6 small nuclear RNA (snRNA) in human cells, a function which does not require the 7SK snRNP (Hasler *et al*, 2020; Hasler *et al*, 2021; Wang *et al*, 2020). Therefore it is possible that this function is conserved in Drosophila, even though we have not observed a general defect in alternative splicing. Another possibility is that Larp7 exerts its additional function through interaction with a closely related 7SK snRNA, which has been recently identified in Drosophila (Nguyen *et al*, 2021). This non-coding RNA was shown to interact with the core partners of the 7SK snRNP suggesting that it holds a similar function as the canonical 7SK snRNA. Double ablation of the 7SK snRNAs should tell whether this condition mimics the molecular phenotypes of the *Larp7* KO.

To conclude, we have shown that the 7SK snRNP function is mostly restricted to neuronal development in vivo, mimicking several clinical defects in human patients, providing novel insights for the importance of this complex in neuron homeostasis.

**Figure EV1:**
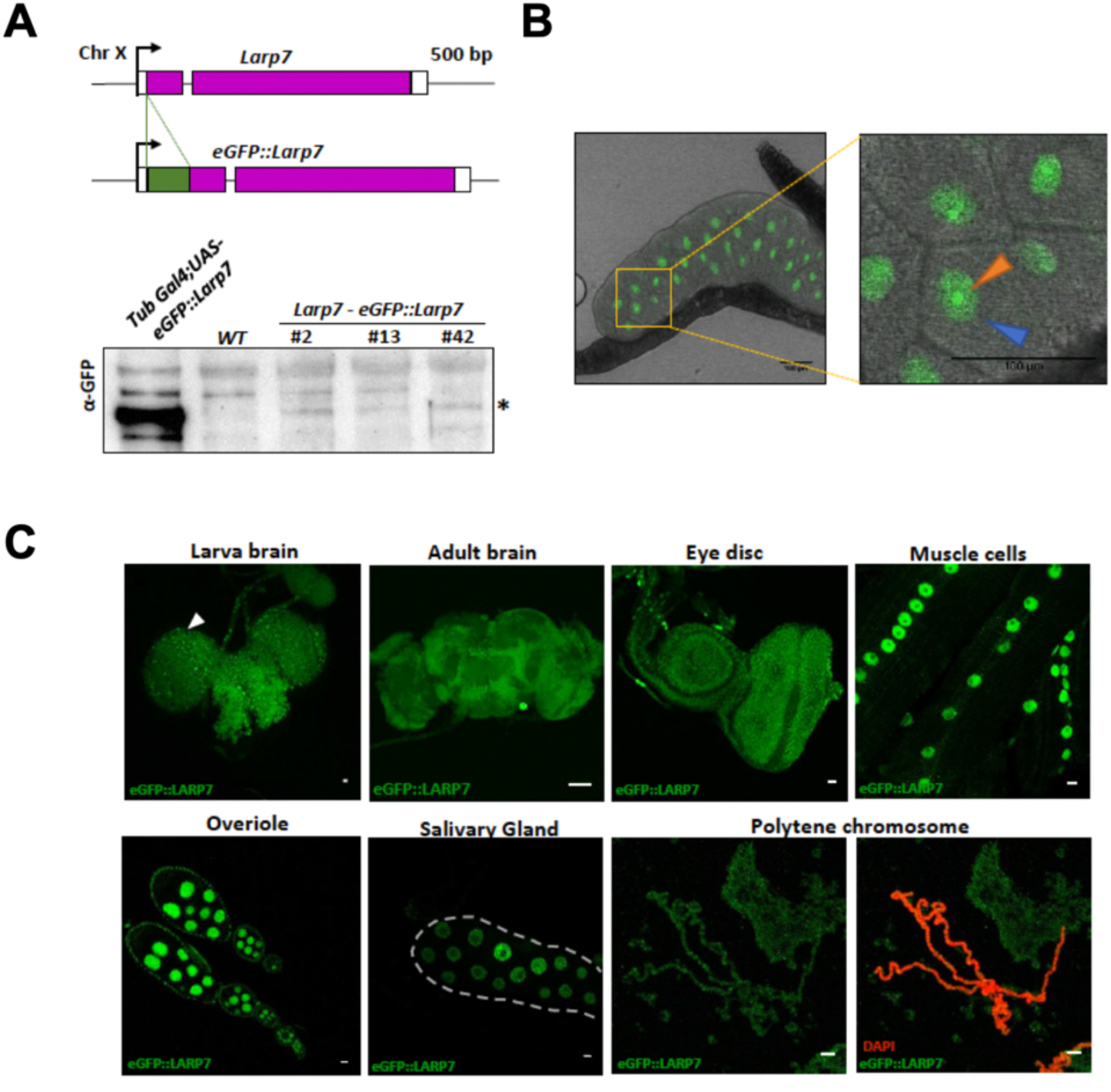
CRISPR/Cas9-mediated GFP tagging of Larp7. **(A)** Top: Scheme of eGFP knock-in at the N-terminal end of the *Larp7* gene. Bottom: Immunoblot using GFP antibody showing expression of eGFP-tagged Larp7 in whole larvae extract. *indicates eGFP-Larp7. Overexpression of UAS-eGFP-Larp7 with *tub*-Gal4 is used for size control. **(B)** Expression of eGFP-Larp7 in salivary gland. Enlarged image shows eGFP-Larp7 localized in the nucleus (blue arrow head) and enriched in the nucleolus (orange arrow head). (**C)** Confocal image of various tissues from larvae and adults depicting eGFP-Larp7 expression.

**Figure EV2:**
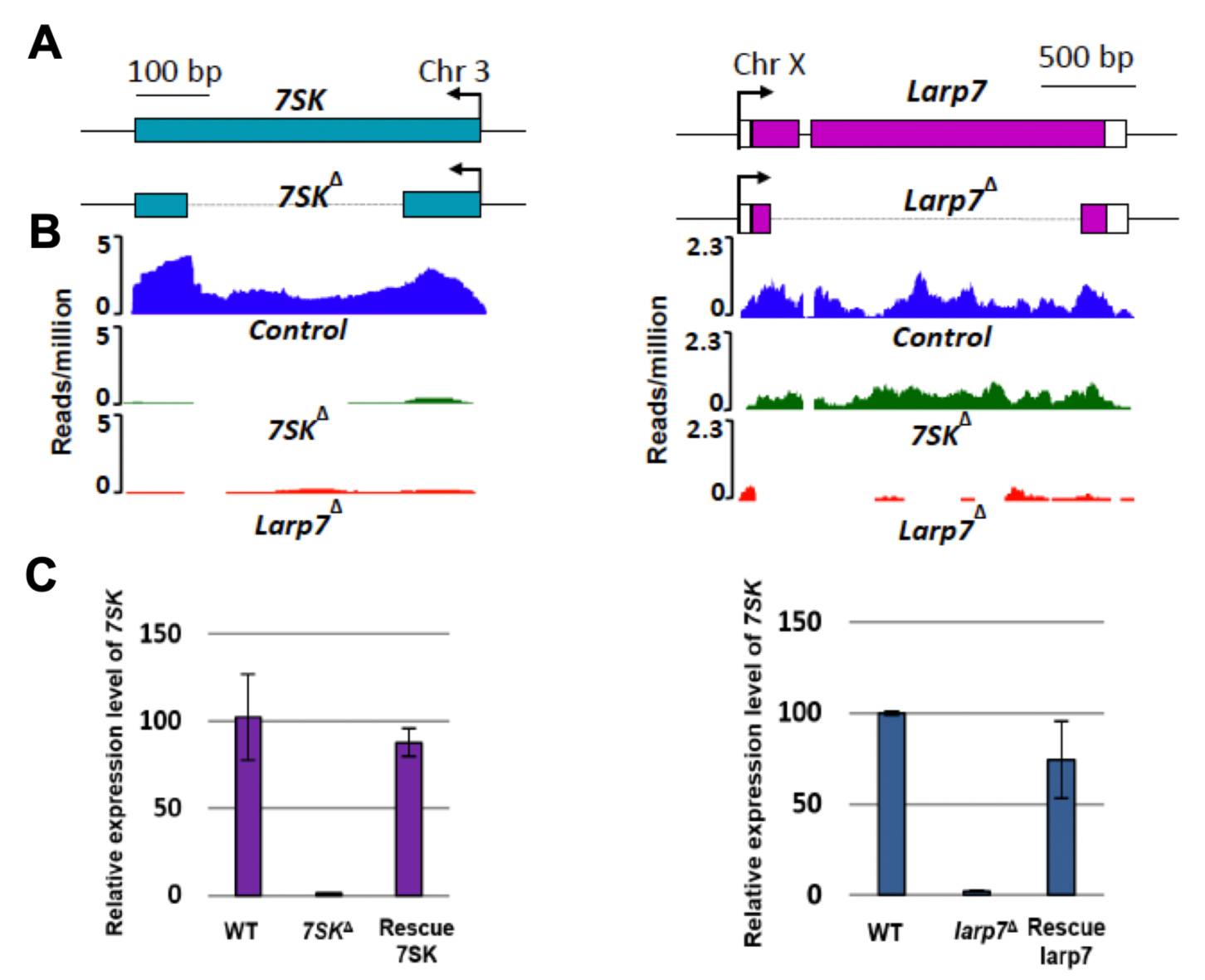
CRISPR/Cas9 mediated knock out of 7SK snRNP. **(A)** Scheme of *7SK* and *Larp7* loci showing the deleted genomic regions in dotted lines. **(B)** UCSC genome browser tracks of RNA-seq data from larvae polyA RNA of control, *7SK* and *Larp7* KO conditions. **(C)** RT-qPCR showing levels of *7SK* RNA in control, *7SK* KO and *7SK* rescued adult flies (left) and control, *Larp7* KO and eGFP-Larp7 rescued adult flies (right). Ten flies were selected for each condition.

**Figure EV3:**
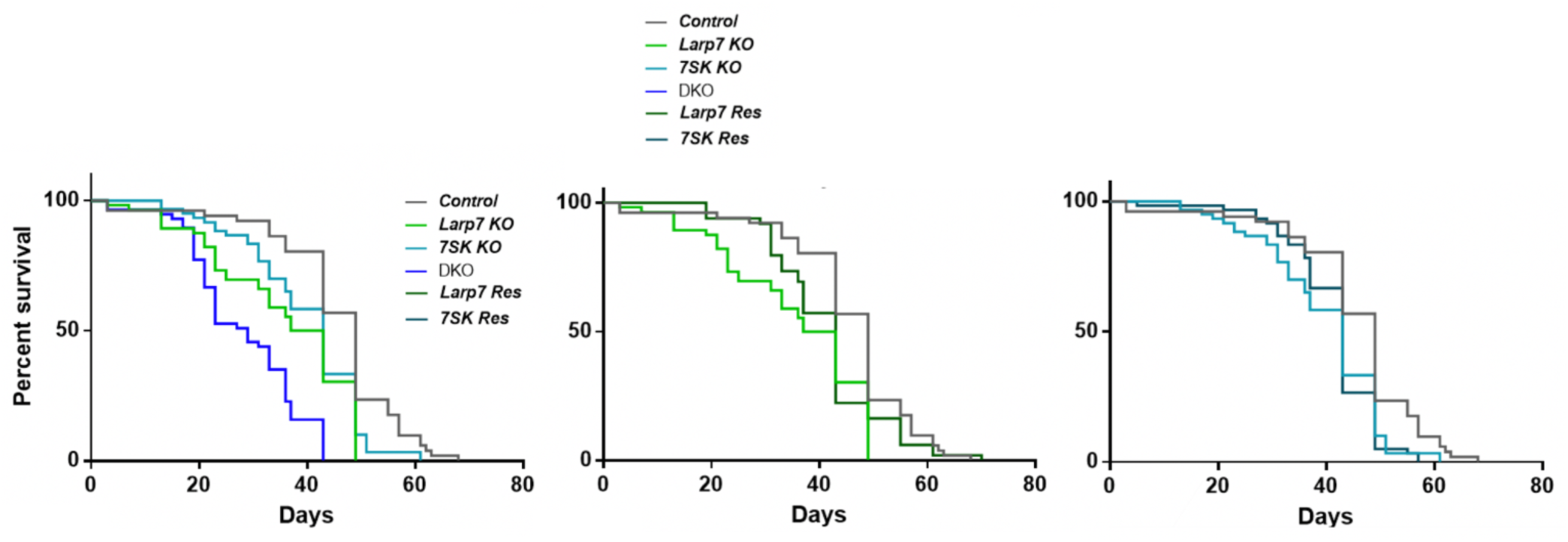
7SK snRNP mutant flies have reduced lifespan. Lifespan of control, *7SK* KO, *Larp7* KO, DKO and rescued flies are shown. Percentage survival is plotted.

**Figure EV4:**
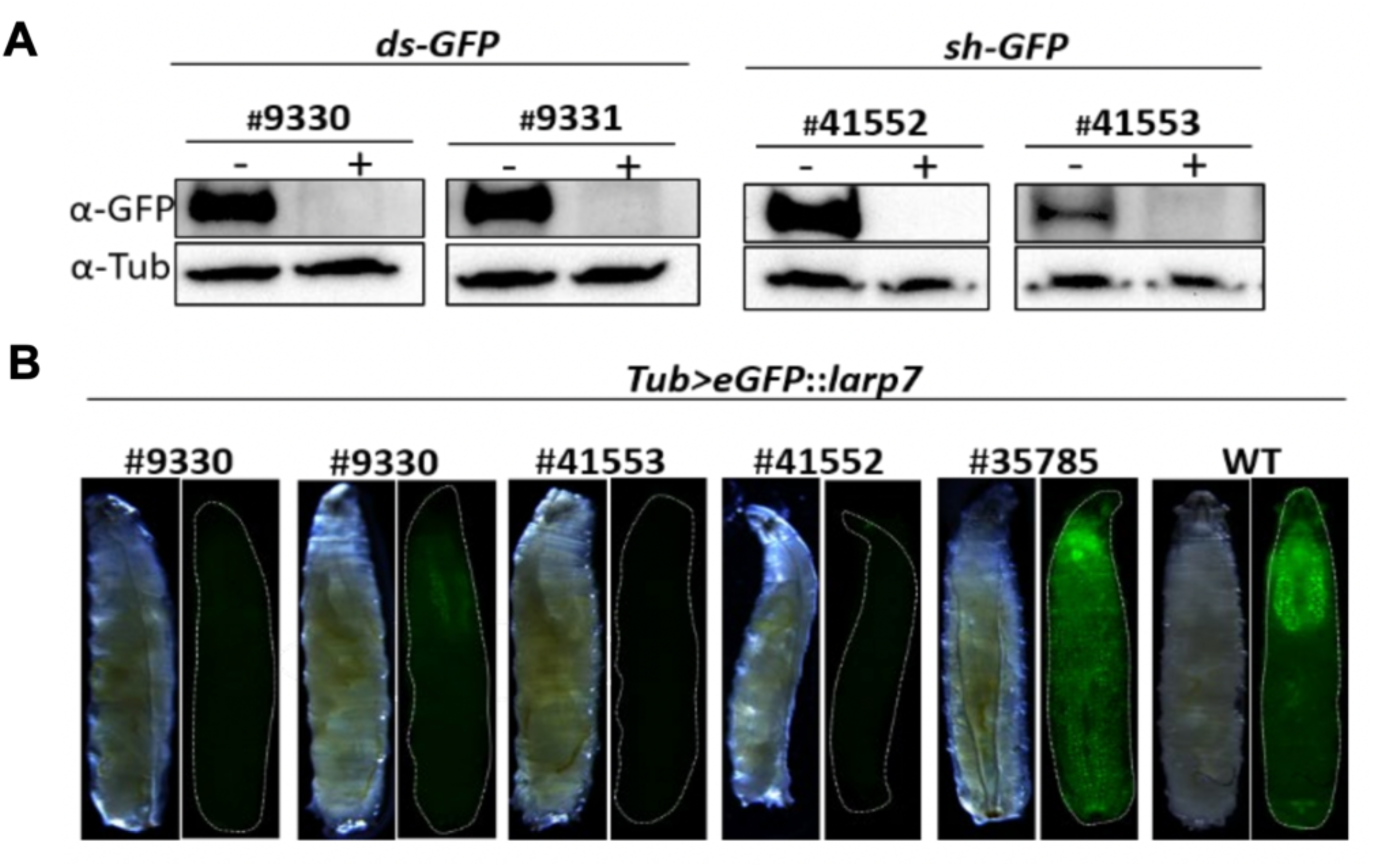
Validation of GFP KD efficiency. **(A)** Immunoblots using anti-GFP antibody and adult head extracts of indicated genotypes. ds-GFP and sh- GFP shows strong reduction in eGFP-Larp7 levels. Anti-Tubulin antibody is used as loading control. **(B)** Bright field and fluorescent images of L3 larvae (*tub*-Gal4>UAS-eGFP::Larp7) ectopically expressing eGFPLarp7. iGFPi lines are indicated. #35785 line contains a transgene driving dsRNA against mCherry, which is used as negative control.

**Figure EV5:**
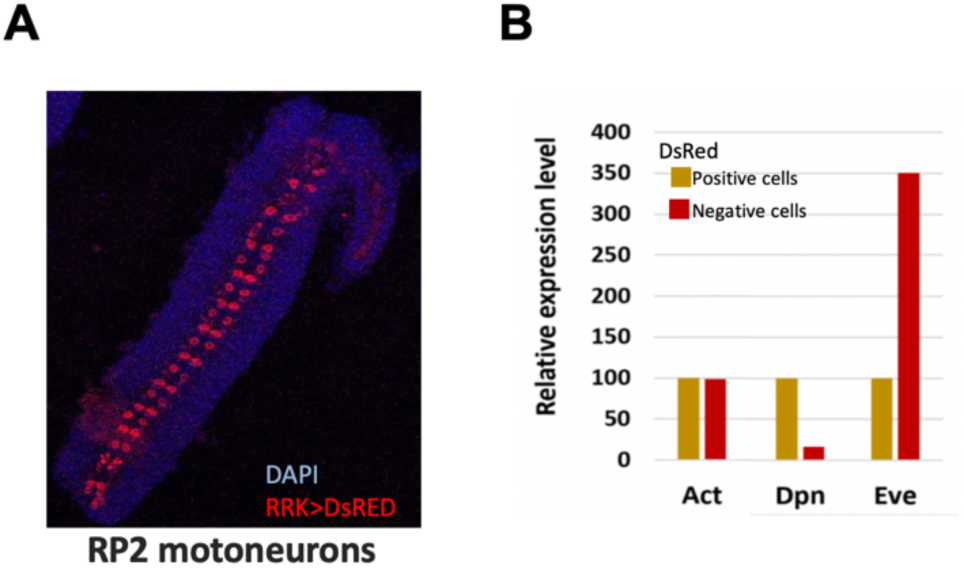
Validation of the cell sorting efficiency. **(A)** Projected confocal images of dissected embryo neuroectoderm expressing nuclear DsRed in RRK positive cells which is used as marker for sorting cells. Nuclei are marked with DAPI (blue). **(B)** RT-qPCR of actin, deadpan (dpn) and even skipped (eve) in sorted DsRed positive and negative cells. DsRed positive cells are enriched with eve confirming the identity of the isolated cells.

## Materials and Methods

### Drosophila Stocks

Flies were maintained at 18°C and 25°C in 60% humidity incubator with 12hrs day – 12hrs light cycle and were fed with standard cornmeal food. The genotypes used in this study are listed in Table 1.

Tools and protocol from the National Institute of Genetics (NIG-Fly) FlyCas9 were used to generate *Larp7* and *7SK* gene deletion and eGFP-*Larp7* endogenous tag. Briefly, for gene deletion, a pair of guide RNAs (gRNA) flanking the gene of interest and for endogenous tagging single gRNA close to the transcription start site (TSS) of *Larp7* was designed using “*Cas9 target finder*”. The gRNA was synthesised (Merck, Germany) and cloned into a pBFv-u6.2/B vector. For gene deletion, the vector was integrated (PhiC31 integrase) into TBX-0002 (*y1 v1 P{nos-phiC31\int.NLS}X; attP40 (II)*) and crossed with CAS-0001 (*y2 cho2 v1; attP40{nos-Cas9}/CyO*). The offspring were individually crossed with respective balancer fly lines, and the parents were screened by PCR (primer sequence in Table 2) for deletion. For endogenous tagging, the donor plasmid was created as follows. Approximately 1000 base pairs (bp) upstream of *Larp7* TSS and 1000 bp downstream of *Larp7* TSS were cloned upstream (BglII) and downstream (KpnI) of eGFP in the p*UAST*-eGFP attB vector (eGFP sequence was cut out). The donor plasmid and gRNA vector pBFv-U6.2 were injected into CAS-0001 embryos. The survived flies were individually crossed with *X* chromosome balancer (*Fm6*,*w)*, after five days or once the offspring L1 larvae hatch, the parent flies were screened by PCR (primer sequence in Table 2) followed by agarose gel electrophoresis. The positive candidates were further verified by PCR-sequencing.

### Gene cloning and generation of transgenic flies

The *Drosophila Larp7* cDNA was synthesised from total RNA of S2R+ cells and cloned in KpnI site of the p*UAST*-eGFP attB vector, eGFP as N-terminal tag. The Human *Larp7* (*hLarp7*) and *Xenopus laevis Larp7* (*xLarp7*) cDNA were synthesised from HeLa cells and *X*. *laevis* embryos, respectively. Further, *hLarp7* and *xLarp7* were cloned in KpnI and NotI site of the pUAST-eGFP-attB vector.

The final clones were verified by sequencing and expression of the eGFP-*hLARP7* and eGFP-*xLarp7* was first validated in S2R+ cells by microscopy and immunoblot. For expression in S2R+ cells, the p*Act*-Gal4 vector was co-transfected to induce the expression of eGFP-*Larp7* under UAS promoter (Brand & Perrimon, 1993). For the *7SK* rescue clone, *7SK* gene and upstream 1500 bps (including transfer RNA: Valine) was amplified by PCR from fly genomic DNA. The amplicon was further cloned into HindIII-EcoRI digested p*UAST*-eGFP attB vector. The cDNA clones were injected into embryos expressing PhiC31 integrase (BestGene Inc., U.S.A) in the genetic background of BDSC #9723 [*y1* w1118; PBac{y+-attP-3B}VK00002]. The transgenic flies were selected based on *white* gene expression form the pUAST vector.

### Single fly genomic DNA

The single fly protocol was used for screening the transgenic flies. Individual flies were collected in 1.5mL centrifuge tubes on ice. Then 50uL of squash buffer (10mM Tris pH8.0, 1mM EDTA, 25mM NaCl) with freshly added 1uL of 10mg/mL Proteinase K was pipetted using a 200uL pipette, and the flies were crushed using the same pipette tip for a few times. Then the sample was incubated at 37°C for 15 min followed by 5 min at 95°C to inactivate Proteinase K. Then spin down the debris at 14,000 RPM for 5 min. Then 5uL of the supernatant was used as a template in a 20uL polymerase chain reaction (PCR).

### Reverse transcription and quantitative real-time PCR

Total RNA were isolated by TRIzolTM reagent. The required amount of total RNA was treated with two units of DNase I (New England Biolabs). For reverse transcription, random oligonucleotides (Sigma, custom made), 200 units of M-MLV reverse transcriptase (Promega), and 18 units of Murine RNase Inhibitor were used according to the manufacturer’s protocol. SYBR Green PCR Master Mix and gene-specific primers (Table 2) and normalisation control primers (*Rpl15*, *Rp49*, and *Act5C*) were used to perform quantitative real-time PCR in ViiA 7 Real-Time PCR system (Applied Biosystems). The relative expression level was calculated using Microsoft Excel, and the student’s t-test was performed for biological triplicates.

### S2 cells KD and transfection

KD of genes in S2R+ cells were done according to the protocol (Rogers & Rogers, 2008) with minor modifications. Genomic DNA or cDNA from flies were used as PCR (OneTaq, NEB) template to amplify the exonic region of genes. The dsRNA was synthesised using MEGAscript™ T7 Transcription Kit as manufactured protocol. The KD treatments were done either once or three cycles for 7 days according to the KD efficiency of individual dsRNA, which was first optimised by titration. KD efficiency was analysed by RT-qPCR (Table 2). For *hLarp7* rescue experiment in S2R+ cells, the cells in six-well plates were treated with dsRNA targeting the 3’UTR of *dLarp7* for five days, three treatments and on the 3rd day, the pUAST rescue construct co-transfected with p*Act5C*-Gal4 in the ratio 4:1 using Effectene reagent as manufactured protocol. The expression of rescue constructs was validated by eGFP fluorescence with a stereo microscope and immunoblot. On Day 5, the eGFP positive cells were sorted (BD FACSAria™ II SORP) into TriZol and RT-qPCR was performed to check the level of *7SK* RNA.

### Microscopy

#### Live imaging

Freshly laid embryos on an apple agar plate were collected on adhesive tape, and chorion membrane was removed using forceps. The ventral side of the embryo was stick to glass bottom of the dish (IBIDI, 35mm high glass bottom dish Cat. No#81158) coated with an adhesive (embryo glue). Then the embryos were covered with halocarbon oil (Voltalef, H3S). Live images were captured for every 15-20 min until the last stage of embryo development using VisiScope (Visitron Spinning disc microscope, Germany), GFP channel. For GFP and mCherry co-localization experiment, stage 15-17 embryos were collected and mounted as described before and imaged using Leica SP5 confocal microscope. The images were visualised and analysed in IMARIS 9.2 Bitplane and ImageJ (Schindelin *et al*, 2012).

#### Expression pattern of Larp7

Larvae or adult flies were dissected and fixed with 4% paraformaldehyde in PBS (137mM NaCl, 2.7mM KCl, 10mM Na2HPO4, 1.8mM KH2PO4, pH 7.4) for 20 min and washed with PBS 10 min three times. Then the samples were mounted in Vectashield with DAPI. Polytene chromosome squashes were prepared as described (Worpenberg *et al*, 2021) and imaged using SP5 microscope.

#### NMJ immunohistochemistry

Crosses of the desired phenotype were set up with a maximum of 20 flies per vial in 25°C and 50% humidity. Wandering L3 larvae were collected and segregated according to the genotype. The larva was dissected on a silicone plate (15cm dish) with cold PBS, and internal organs were removed. Then fixed with 4% Paraformaldehyde (PFA) in PBS for 45 min and washed in PBS for one hour. The preparations were transferred to 35mm dish with silicone surface and washed six times with PBST (PBS+0.5% Triton-X) for 30 min each wash. Primary antibody anti-Syt 1:200 (3H2 2D7, Developmental Studies Hybridoma Bank, DSHB) in 0.3% PBST was added and incubated overnight at 4°C. The antibody was removed and washed eight times with PBS for each 30 min. The secondary antibody, anti-mouse-Alexa 488 or anti-mouse-Cy3 (1:500) and Horseradish Peroxidase (HRP)-conjugated TRITC [Tetramethylrhodamine-5-(and 6)-isothiocyanate] (1:1000) was added and incubated in 4°C, overnight. Finally, washed six times with PBS and mounted with Vectashield. NMJ from muscle four was imaged from segment A2 and A3 for NMJ analysis. Nerves were identified using anti-HRP-TRITC staining and anti-Syt (DSHB) to mark the synapses. Z stack for the whole NMJ was imaged with 0.4uM thickness. The images were further analysed using IMARIS 8 image analysis software.

#### Immunostaining

Embryos laid overnight by the desired genotype flies were collected on an apple agar plate. Then dechorinate with 50% bleach (sodium hypochlorite) for 2 min and washed with deionised water on 100µM cell strainer. The embryos were next transferred to 1.5mL centrifuge tubes containing (1:1) 4% Paraformaldehyde: n-Heptane and placed on a shaker for 20 min at 500 RPM in room temperature. Then spun down at 500 G and the lower phase (fix) was removed, and an equal volume of methanol was added and mixed for one minute. Then washed with methanol for two more times and then stored in -80°C. The fixed embryos were rehydrated with PBS and incubated in primary antibody in PBSB (PBS + 5% Donkey serum) overnight at 4°C. The embryos were washed with PBT, 30 min, four times and secondary antibody [Alexa 488, Cy5 and Cy3 (1:1000)] was added and incubated overnight at 4°C. Finally, the embryos were washed with PBT and mounted with Vectashield containing DAPI. The embryos were imaged using SP5 confocal microscope.

#### Crawling assay

Properly staged wandering L3 larvae were collected and briefly washed with water. Then the larva was placed on the centre of the 15cm agar plate (1%). The locomotion was observed for 2 minutes after the recovery time for up to 1 min. The number of peristaltic movements was counted manually by observing the larvae or analysed recorded videos in IMARIS software.

#### Lifespan assay

Five days old, 20 flies were collected for each condition and placed in a vial containing apple agar and a layer of dry yeast and maintained at 25° C, 55% humidity with 12hrs dark-light cycle. Flies were transferred to new food every third day and recorded the number of dead flies. The observation was done until all the flies died in all conditions. Survival graph analysis form PRISM 8 was used to perform statistical analysis. Oasis online tool (Yang *et al*, 2011) was used to plot the mortality rate between the samples.

#### Bouton calculation

The NMJ images were acquired as described before. The images were visualized and analyzed in IMARS 8 as follows. The surface application was used to mark all the boutons in the NMJ muscle 4. Then the number of boutons were counted manually and recorded in GraphPad (PRISM 8), and statistical tests were performed according to the distribution of the data.

#### Immunoblot

About five heads of adult flies were squashed in 50ul of 2xLDS buffer (NuPAGE), then incubated at 90°C for 10 min, cool down to room temperature and spun at 14,000 RPM for 10 min. 20ul of the sample was loaded in 10% SDS gel and resolved. The proteins were transferred onto PVDF membrane and blocked with 5% milk. The membrane was incubated with primary antibody overnight and with secondary antibody for 2 hours and imaged using Pico or Femto reagents (ThermoFisher). The images were captured using a BioRad GelDoc system and quantified using ImageJ software.

#### RNAi in larvae

To KD *Larp7* in flies, we used endogenously tagged eGFP-*Larp7* flies. First eGFP-*Larp7* flies were recombined with UAS-*Dicer* (*X* chromosome) to increase the efficiency of KD. Then sh-*GFP* (#41552) in the second chromosome was recombined to get eGFP-*Larp7*, UAS-*Dicer*; sh-*GFP*. The KD efficiency of all available GFP RNAi lines was analysed by immunoblot and imaging. sh-RNA line #41552 and #41553 showed the strongest depletion of GFP, but #41553 was less fertile than #41552. Due to this, we used #41552 to for all the KD experiments. Then virgin female of eGFP-*Larp7*, UAS-*Dicer*; sh-GFP line crossed with different Gal4 drivers and maintained the crosses in 25°C. The crawling assay was performed with F1 male L3 larvae, for *Cdk9* knockdown #34982 line was used since it shows the highest KD efficiency (Akhtar *et al*., 2019). *dsCdk9* (3rd Chr) was recombined with *RRK*-Gal4 line and the recombinants were screened by PCR (Table 2) and sequencing.

#### RNA sequencing

The wild-type control, *Larp7* KO and *7SK* KO flies were recombined with UAS-nuclear DsRed (BDSC #8546, w[1118]; P{w[+mC]=UAS-RedStinger}4/CyO). Then these flies were crossed with *RN2*-Gal4 (BDSC #7470) or *RRK*-Gal4 (BDSC #42739), and the expression pattern of DsRed was verified by confocal imaging of embryos and L3 larvae. Then *Larp7* KO and *7SK* KO flies were recombined with *RN2*-Gal4 line. *RRK*-Gal4 line could not be recombined with *7SK* mutants. Nevertheless, we did not see any difference in expression pattern between these two lines, therefore, *RN2*-Gal4 line was used in this experiment. The crosses were maintained in a plastic flask with apple agar plate coated with yeast. After two pre-laying every two hours, embryos were collected for 2 hours on an apple agar plate. The staged (stage 15-16) embryos were collected and processed as described (Perrimon *et al*, 2011) with minor modifications. The embryos were decorated with 50% bleach (sodium hypochlorite) for 2 min and washed thoroughly with deionised water in 100µM cell strainer. Then transferred to 250 uL Schneider’s *Drosophila* Medium (Gibco) and dissociated using Dounce homogenizer with loose pestle 15 times on the ice. Then the dissociated cells were filtered through the 35µM cell strainer and collected in FACS compatible 5mL tube on ice. To the dissociated cells, DAPI was added to eliminate dead cells during sorting and immediately analyzed in FACS. The purity of the sorting was analysed visually by microscope. Further, RNA was isolated from about 250 DsRed cells, and RT-qPCR was performed to check the marker DsRed and *eve* transcripts. For RNA sequencing, the DsRed positive live single cells were sorted in 250uL TriZol reagent at 4°C. The sorted cells were frozen on dry ice and stored at -80°C. Then the total RNA was isolated according to the TriZol manufacture’s protocol. The quality of the RNA was analysed using Agilent RNA 6000 Pico assay in Agilent 2100 Bioanalyzer and quantified using NanoDrop 3300. Equal concentration of RNA from each biological triplicate were used in Smart-Seq2 Illumina protocol to generate next-generation sequencing libraries.

#### Raw-data Processing

Libraries were sequenced on an Illumina NextSeq 500 in single read mode with a read length of 84 bp on a single flowcell and sorted according to their index using bcl2fastq (v2.19). Individual samples were first mapped against an rRNA reference to remove ribosomal reads and then against BDGP6 (Ensembl release 90). The mapper used was STAR (Dobin *et al*, 2013). Gene counts were derived using featureCounts (Liao *et al*, 2013).

#### RNA-Seq Analysis

Differential expression analysis was performed using Bioconductor/DESeq2 (Huber *et al*, 2015; Love *et al*, 2014). Genes were deemed sign. diff. regulated with and FDR below 5%.

#### GC content

The GC content around the TSS/Metagene are based on the differentially regulated genes derived from the DESeq2 analysis. For each gene, the TSS with the highest coverage and the associated transcript was chosen for display. GC content around the TSS was calculated using R/Bioconductor packages ((Huber *et al*., 2015; Lawrence *et al*, 2013) (R Core Team 2018) and plotted using ggplot2 (Wickham, 2016). GC content on the metagene profiles were based on GC content per bp retrieved from the genome using bedtools (Quinlan & Hall, 2010) and kentUtils (Kent, n.d.). Then the data were processed further using deeptools2 (Ramirez *et al*, 2016) and plotted using ggplot2 (Wickham 2016).

#### Gene Length

Length of genes and introns were derived from Ensembl release 90 using the GenomicFeatures and GenomicRanges packages (Lawrence *et al*., 2013) and the plotted using ggplot2 (Wickham 2016).

#### Motif occurrences

Motif occurrences were quantified using the genomation package (Akalin *et al*, 2015). A hit was assumed if the sequence had a minimal score of 0.8. Meta profiles and non density heatmaps around the TSS of motif occurrences were plotted using genomation. Density heatmaps were produced using customs scripts recreating an output equivalent to the density heatmaps produced by seqPattern (https://rdrr.io/bioc/seqPattern/) and plotted using genomation. For PauseButton and GAGA consensus motifs were taken from (Hendrix *et al*, 2008) and used for motif occurence quantification. For INR, DPE, DRE and TATA motifs, position frequency matrices were retrieved from (Rach *et al*, 2009) and used for the quantification.

### Data Availability Section

All data needed to evaluate the conclusions in the paper are present in the paper and/or Supplemental Materials. Additional data related to this paper may be requested from the authors.

### Disclosure and competing interests statement

The authors declare no competing interests.

## Acknowledgements

We thank the Bloomington and Vienna stock Center for fly lines and the *Drosophila* Genomics Resource Center at Indiana University for plasmids and cell lines; We indebted to BestGene for injections. We thank all the members of the Roignant lab for helpful discussion. Support by the IMB Genomics Core Facility and the use of its NextSeq500 (funded by the Deutsche Forschungsgemeinschaft (DFG, German Research Foundation) – INST 247/870-1 FUGG) is gratefully acknowledged. Research in the laboratory of J.-Y.R. is supported by University of Lausanne, funds from the Swiss National Science Foundation (310030_197906) and the Deutsche Forschungsgemeinschaft (project number 439669440 TRR319 RMaP TP B01)

## Notes

### Competing Interest Statement

The authors have declared no competing interest.

